# Extremely low sample size allows age and growth estimation in a rare and threatened shark

**DOI:** 10.1101/2022.09.26.509619

**Authors:** Peter M. Kyne, Jonathan J. Smart, Grant Johnson

## Abstract

Understanding life history parameters is key to assessing biological productivity, extinction risk, and informing the management of exploited fish populations. Age-and-growth analyses in chondrichthyan fishes (sharks, rays, and ghost sharks) is primarily undertaken through counting band pairs laid down in vertebrae. For rare, threatened, and protected species such as river sharks (family Carcharhinidae; genus *Glyphis*) of northern Australia, obtaining sufficient samples of vertebrae may not be possible. Here we use a very sample size, selective size-class sampling, and back-calculation techniques to provide age and growth data on the Speartooth Shark *Glyphis glyphis* from which comprehensive sampling is not possible. Ten individuals were sampled from the Adelaide River, Northern Territory, Australia. Length-at-age models were applied to the observed and back-calculated data with the sexes combined due to the small sample size and growth estimated using a multi-model framework. Band pair counts produced age estimates of 0–11 years. Most model parameter estimates for length-at-birth (*L*_*0*_) and asymptotic length (*L*_*∞*_) were biologically plausible. The model averaged parameters for the observed data were 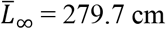 total length (TL) and 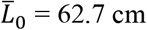 TL, and for back-calculated data were 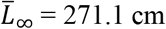 TL and 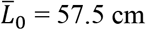 TL. Overall, the parameter standard errors and model residual standard errors were lower for the back-calculated data due to the addition of interpolated data. Analysed samples were restricted to juveniles and sub-adults as adult *G. glyphis* have not been encountered in the Northern Territory. The ageing results suggest an age-at-maturity of >12 years for this species. The lack of mature individuals in the sample means that this analysis should be considered as a partial growth curve with length-at-age estimates that are valid over the available age range. The results presented here provide the first age and growth estimation for river sharks.

## Introduction

Understanding species life history parameters is key to assessing biological productivity, extinction risk, and informing the management of exploited populations. Biologically productive species are able to withstand higher exploitation levels and recover more quickly if populations are over-exploited or depleted (Musick 1999). The collection of life history data most often requires lethal sampling to understand key aspects such as age and growth and reproductive biology (Heupel and Simpfendorfer 2010). Age, fecundity, reproductive periodicity, and mortality estimates (the latter of which can be calculated from growth data) are required to calculate demographic parameters. This includes productivity measures such as intrinsic rates of population increase (e.g., Smith et al. 1998). Age data are also required to calculate generation length, a key demographic parameter when assessing species’ extinction risk within the framework of the IUCN Red List of Threatened Species (IUCN Standards and Petitions Committee 2019).

Unsustainable exploitation and trade are driving a heightened level of extinction risk among the world’s chondrichthyan fishes (sharks, rays, and ghost sharks; hereafter ‘sharks’). About a third of all sharks are estimated to be threatened with extinction (IUCN 2022). Amongst non-marine sharks, a highly specialised group of relatively low diversity (Grant et al. 2019), euryhaline species are particularly at risk. These species can tolerate the range of salinities from marine to freshwaters but occupy restricted habitats within riverine and estuarine environments (Grant et al. 2019, Kyne and Lucifora 2022). Of the 10 euryhaline sharks globally, six are threatened including three Critically Endangered species (IUCN 2022, Kyne and Lucifora 2022).

The river sharks (family Carcharhinidae; genus *Glyphis*) comprise three threatened euryhaline species of the Indo-West Pacific. Two species (Northern River Shark *G. garricki* and *G. glyphis*) have patchy restricted geographic ranges in northern Australia and southern Papua New Guinea (Feutry et al. 2020, Kyne et al. 2021). They display a high degree of population structuring (Feutry et al. 2017, Feutry et al. 2020, Kyne et al. 2021) but their basic life history (e.g., age and growth, reproductive biology) is poorly known (Pillans et al. 2009). Both species are protected as a result of their threatened listings under Australia’s national environmental legislation.

This protected status precludes the assessment of life history parameters required for demographic analyses and management purposes. Ideally, shark ageing studies utilise a large sample size to obtain accurate age and growth parameters and to be able to assess these parameters separately for each sex. Large sample sizes are not always feasible for fish species with low encounter rates with fisheries (e.g., due to rarity or catchability) or for species of conservation concern. Here we use a very sample size, selective size-class sampling, and back-calculation techniques to provide age and growth data on the rare, threatened, and protected *G. glyphis* from which comprehensive sampling is not viable. We provide plausible age and growth parameters using the smallest sample size of any shark study and provide the first such parameters for river sharks.

## Methods

### Sample collection

Samples were collected from the middle to lower reaches of the Adelaide River, Northern Territory, Australia. Sharks were caught between 08 December 2015 and 23 November 2016 from a boat using rod-and-line on fresh teleost fish bait. To obtain a wide range of specimens for vertebral ageing, sharks were targeted at ∼20 cm intervals between size-at-birth (50–65 cm total length (TL); Pillans et al. 2009) and the maximum recorded size in the Northern Territory of ∼190 cm TL (P.M. Kyne, unpubl. data). Upon capture, sharks were lifted into a sampling tub, identified, measured, weighed, and photographed. If the individual *G. glyphis* met the desired size-class for vertebral sampling, the shark was immediately euthanised using a saturation bath of MS-222. Jaws of all processed sharks have been deposited in the Museum and Art Gallery of the Northern Territory in Darwin. All other elasmobranchs, including additional *G. glyphis* outside desired size classes, were processed for tagging and tissue sampling as part of a larger project and released at the site of capture.

Maturity was assessed in each individual shark by examining the state of external clasper calcification in males, and the state of internal reproductive tract development in females. Uncalcified claspers indicated an immature male. Undeveloped internal reproductive tract indicated an immature female; developing internal reproductive tract including oviducal gland width >10 mm indicated a sub-adult female.

All sampling adhered to permitting and ethical requirements and was undertaken under Northern Territory of Australia *Fisheries Act* Special Permit No 2014-2015/S17/3364, and Charles Darwin University Animal Ethics Committee Project Application and Permit Approval A11041.

### Vertebrae sectioning

Vertebrae were processed following protocols described by Cailliet and Goldman (2004). Once transported to the laboratory, the vertebrae were defrosted, and remaining muscle tissue was removed using a scalpel. Individual vertebral centra were then separated and soaked in a 4% sodium hypochlorite solution for 30 mins to remove any remaining tissue. Centra were then dried in an oven at 60°C for 24 hours. A low speed circular saw with two diamond-tipped blades (Beuhler, Illinois, USA) was used to section individual centra. These longitudinal sections were made through the centrum focus at a thickness of 400 µm. Sections were mounted onto microscope slides using Crystal Bond adhesive (SPI supplies, Pennsylvania, USA).

### Age determination

Ages of individual centra were estimated by counting the translucent and opaque band pairs in the *corpus calcareum* under a microscope using transmitted light (Cailliet and Goldman 2004). A change in angle of the *corpus calcareum* was interpreted as a transition from pre-to post-natal growth and marked an age of zero (Fig. 1). Each subsequent growth band pair was assumed to be one year of growth. Annual growth band deposition could not be validated in this study due to the sampling limitations.

**Figure 1.**
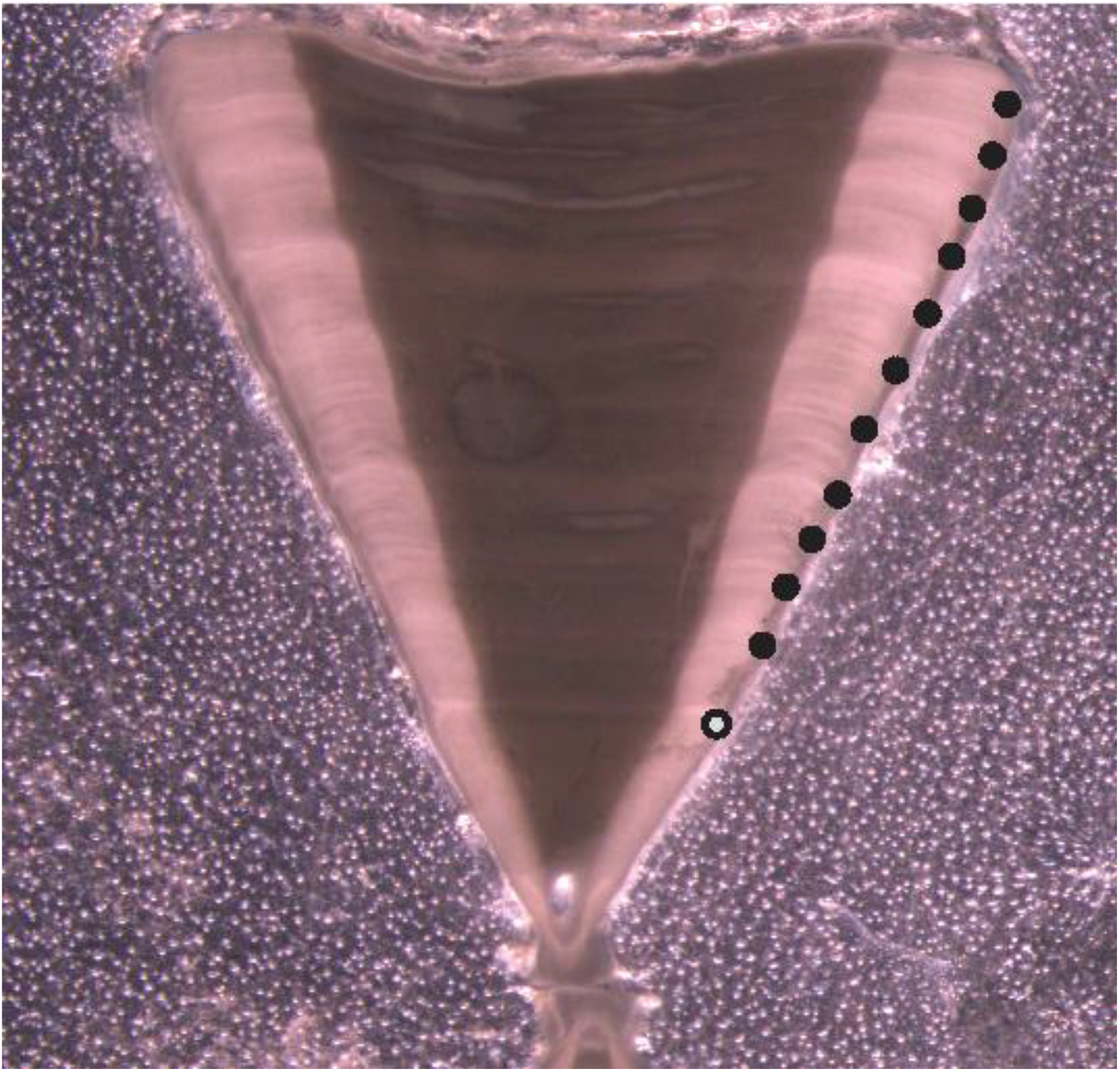
Photograph of a sectioned vertebral centrum from a 181.5 cm total length female *Glyphis glyphis* with 11 visible growth band pairs (solid black circles). Age zero (birth mark) is denoted by the grey circle with a black outline.

Growth band pairs were counted by two independent readers to reduce age estimation bias (Campana et al. 1995, Cailliet and Goldman 2004). When counts differed between readers, the samples were re-examined until a consensus age was reached. If no consensus age was reached, that centrum was removed from analysis. Percent agreement (PA) and percent agreement ± 1 year (PA ± 1 year) were calculated between growth band reads (Cailliet and Goldman 2004). However, standard inter-reader precision statistics such as Bowker’s test of symmetry (Bowker 1948, Evans and Hoenig 1982), average percent error (APE), and Chang’s coefficient of variation (CV) (Chang 1982) were precluded by the small sample size (Smart et al. 2013).

### Back-calculation

Back-calculation techniques were used to supplement the low sample size with interpolated data (Francis 1990). Centra were photographed using a compound video microscope and the distances between growth band pairs were measured using image analysis software (Image Pro Plus version 6.2 for Windows, Media Cybernetics, 2002). The centrum radius (*CR*) was measured as a straight line from the focus to the centrum edge. The birth mark and each growth band pair were measured along this line as the distance from the focus to the nearest 1 µm. A Dahl Lea direct proportions back-calculation technique was applied (Carlander 1969):

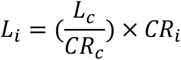

where *L*_*i*_ = length at growth band pair ‘*i*’, *L*_*c*_ = length at capture, *CR*_*c*_ =centrum radius at capture and *CR*_*i*_ = centrum radius at growth band pair ‘*i*’. An assumption of the Dahl Lea direct proportions method is that there is a linear relationship between *L*_*c*_ and *CR*_*c*_. This was tested by performing a linear regression between these two measurements. A Fraser Lee size-at-birth modified back-calculation approach was also applied (Campana 1990). However, visual inspection showed that the Dahl Lea direct proportions approach provided improved estimates and thus was used for all further analyses.

### Growth modelling

Length-at-age models were applied to the observed and back-calculated data with the sexes combined due to the small sample size (Smart et al. 2013). Growth was estimated using a multi-model framework (Smart et al. 2016) that included three candidate growth functions *a priori*: von Bertalanffy growth function (VBGF) (von Bertalanffy 1938), Gompertz function (Ricker 1975), and logistic function (Ricker 1979) (Table 1). Model selection was determined using Akaike’s information criterion (Akaike 1973) with a small sample size adjusted bias correction (*AICc*) (Zhu et al. 2009). Parameterisations that included length-at-birth (*L*_*0*_) and asymptotic length (*L*_*∞*_) parameters were used for all three candidate models (Table 1). These parameterisations are recommended for sharks as they provide more biologically relevant information and can be compared directly between candidate models (Cailliet et al. 2006, Smart et al. 2016). Best fit parameter estimates were determined for all three candidate models using the ‘nls’ function in the ‘R’ program environment (R Core Team 2013). *AICc* was also calculated in the ‘R’ program environment as:

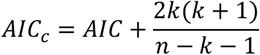

where *AIC = nlog*(σ^2^) + 2*k, k* is the total number of parameters +1 for variance (σ^2^) and *n* is the sample size. The model with the lowest *AICc* value (*AIC*_*min*_) had the best fit to the data and was thus identified as the most appropriate of the candidate models. The remaining models were ranked using the *AIC* difference (*Δ*) which was calculated for each model (*i* = 1–3) as:

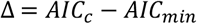

**Table 1.** Model equations of the three *a priori* growth functions used to estimate length-at-age.

| Growth function | Equation | Reference |
| --- | --- | --- |
| von Bertalanffy growth function (VBGF) | $L_t = L_0 + (L_\infty - L_0) (1 - \exp(-kt))$ | von Bertalanffy (1938) |
| Gompertz function | $L_t = L_0 \exp \left( \ln \left( \frac{L_\infty}{L_0} \right) (1 - \exp(-g_{gom}t)) \right)$ | Ricker (1975) |
| logistic function | $L_t = \frac{L_\infty L_0 (\exp(g_{log}t))}{L_\infty + L_0 (\exp(g_{log}t) - 1)}$ | Ricker (1979) |

where $L_t$ is length-at-age $t$ , $L_0$ is length-at-age 0, $L_\infty$ is asymptotic length, $k$ , $g_{log}$ and $g_{gom}$ are the different growth coefficients of the respective models (which are incomparable).

Models with *Δ* of 0–2 had the highest support while models with *Δ* of 2–10 had considerably less support and models with *Δ* of >10 had little or no support (Burnham and Anderson 2001). *AIC* weights (*w*) represent the probability of choosing the correct model from the set of candidate models and were calculated for each model (*i* = 1–3) as:

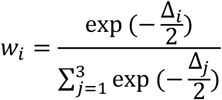

Multi-model inference (MMI) is recommended when no model candidate is the outright best model for the data (*w*_*i*_ > 0.9) (Katsanevakis and Maravelias 2008). Therefore, model averaged length-at-age estimates, parameters, and standard errors were calculated when candidate models had similar *w*. Only *L*_*∞*_ and *L*_*0*_ were comparable between the three model candidates as the three growth completion parameters (*k, g*_*log*_ and *g*_*Gom*_) are incomparable between them. Therefore, a model averaged *L*_*∞*_ was calculated as:

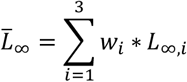

where 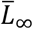 was the model averaged asymptotic length (Burnham and Anderson 2002; Katsanevakis 2006). The unconditional standard error of 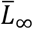 was estimated as:

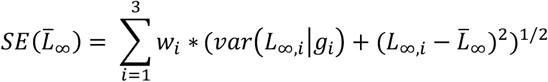

where *var*(*L*_*∞,i*_ | *g*_*i*_) is the variance of parameter *L*_*∞*_ of model *g*_*i*_ (Katsanevakis and Maravelias 2008). A model averaged estimate and standard error of *L*_*0*_ were calculated using the same equations.

## Results

Retained sharks were collected at depths of 4.3–10.5 m (baits were set on the river bottom), water temperatures of 30.9–31.9°C, surface salinities of 4.04–19.06, and turbidities of 25–279 NTU, and consisted of 3 male *G. glyphis* (101.5–153.2 cm TL) and 7 female *G. glyphis* (59.8–189.0 cm TL). All individuals were immature although the largest three females (161–189 cm TL) were classed as sub-adults. The length-weight relationship for both sexes was:

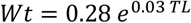

where *Wt* is weight in kg (Fig. 2).

**Figure 2.**
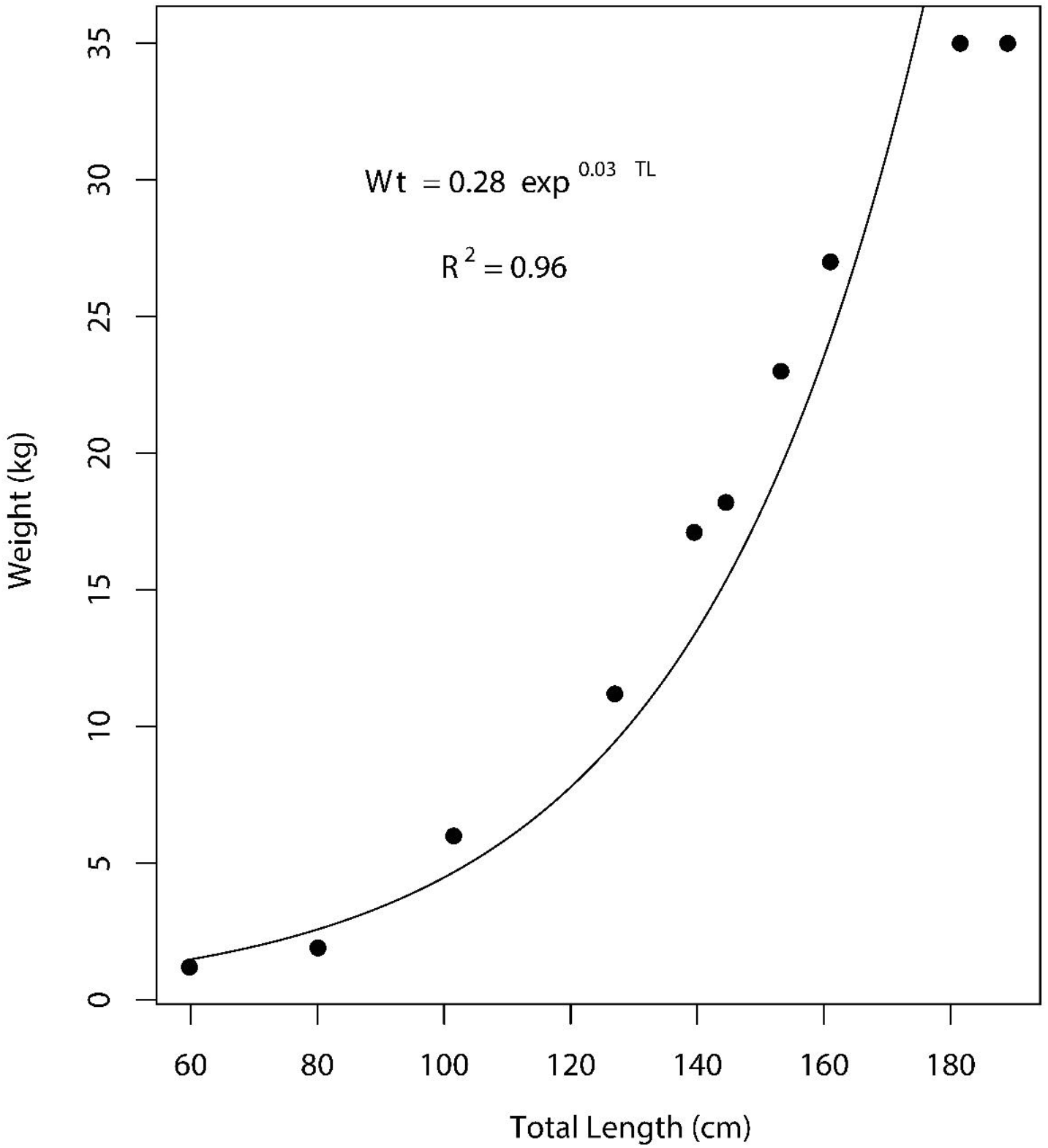
Length-weight relationship for *Glyphis glyphis* (*Wt* = 0.28 exp^0.03 TL^, R^2^ = 0.96, *F*_*1,8*_, p < 0.001).

Male ages were 4, 5, and 8 years while females ages ranged from 0–11 years. The growth band pairs were visible in all centra and easy to identify for most individuals (Fig. 1). This lead to high reader agreement as PA was 46 % and the PA ± 1 year was 82 %. Final ages were agreed upon through consensus reads so no individuals were removed from the analysis. A linear relationship was determined between *L*_*c*_ and *CR*_*c*_ (Fig. 3). Therefore, the Dahl Lea direct proportions technique was appropriate for *G. glyphis*.

**Figure 3.**
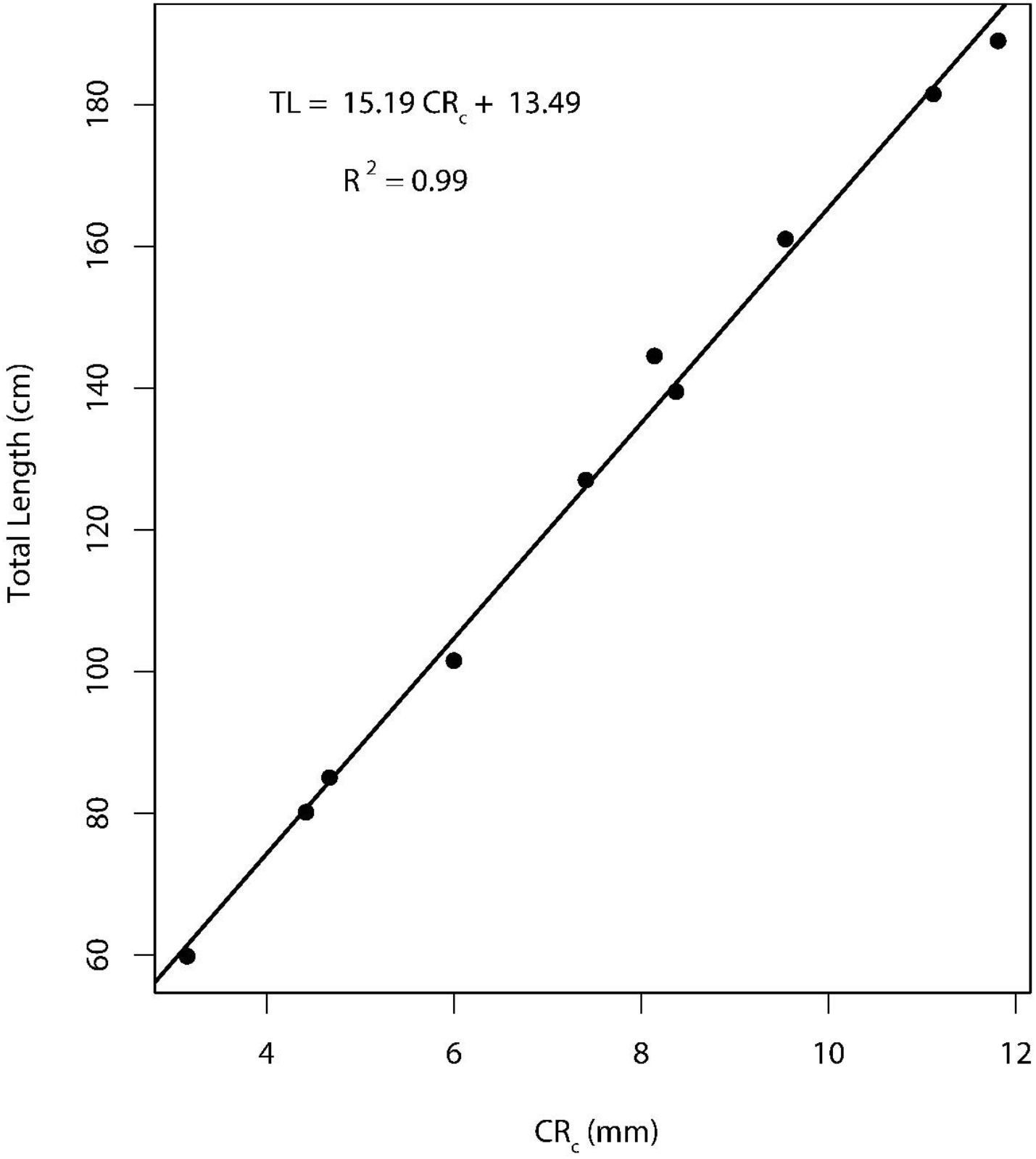
Relationship between centrum radius (*CR*_*c*_) and total length (TL) for *Glyphis glyphis* (TL = 15.19 *CR*_*c*_ + 13.49, R^2^ = 0.99, *F*_*1,8*_, p < 0.001).

Neither the observed data nor the back-calculated data had a candidate growth model where *w*_*i*_ > 0.9 (Table 2). Therefore, MMI was performed for both data sets. Equal *w* occurred for all three candidate models for the observed data despite large differences in *L*_*∞*_ (Table 1). The VBGF provided the largest *L*_*∞*_ (366.3 cm TL) which is larger than the estimated maximum size for the species (∼260 cm TL; White et al. 2015). However, the logistic and Gompertz models produced *L*_*∞*_ estimates that were more biologically plausible (217.8 and 248 cm TL, respectively) (Table 2). All three candidate models produced *L*_*0*_ estimates that were within the range of empirical length-at-birth estimates (50–65 cm TL; Pillans et al. 2009) (Table 2). These *L*_*0*_ estimates had far lower standard errors than the *L*_*∞*_ estimates. This is a common occurrence when older individuals are not included in the sample as greater uncertainty occurs around the *L*_*∞*_ parameter (Smart et al. 2017). The model averaged parameters for the observed data were 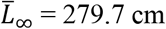 TL and 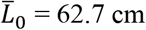 TL (Table 2).

**Table 2.**
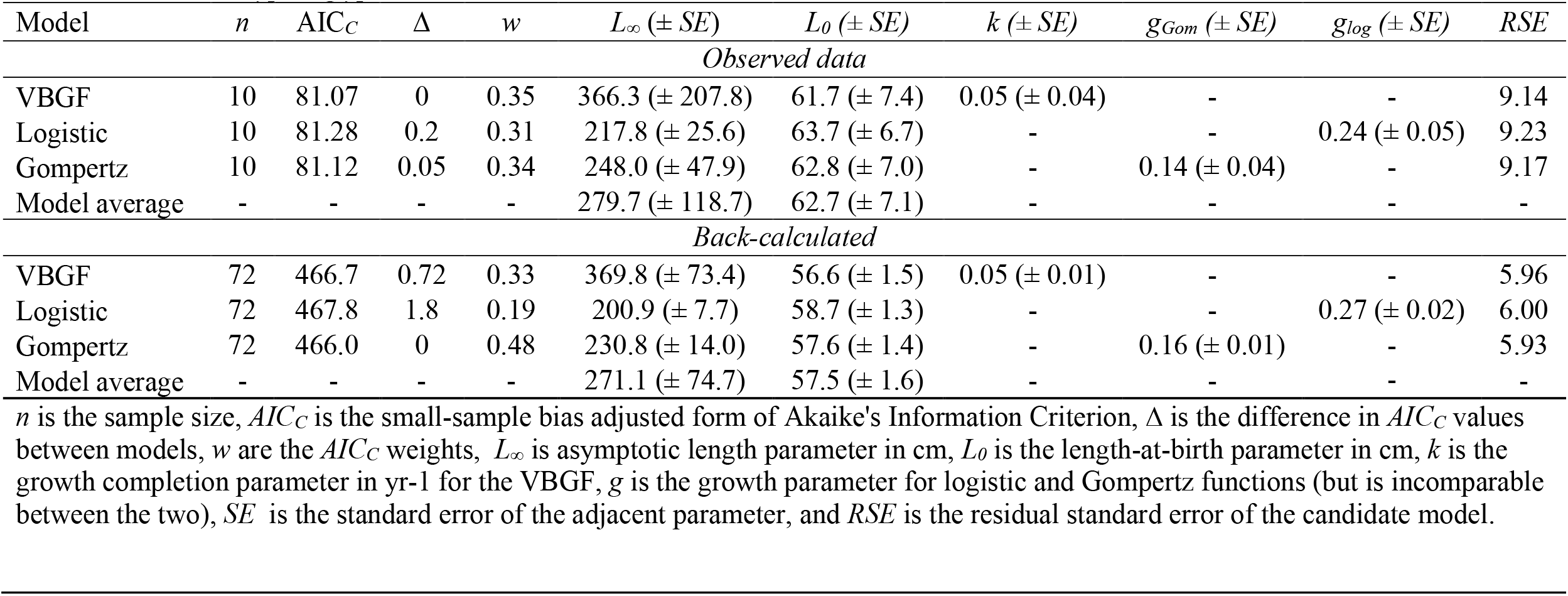
Summary of candidate growth model parameters, MMI parameters and *AIC*_*c*_ results for the observed length-at-age and back-calculated data for *Glyphis glyphis* with sexes combined.

| Model | $n$ | $AIC_C$ | $\Delta$ | $w$ | $L_\infty (\pm SE)$ | $L_0 (\pm SE)$ | $k (\pm SE)$ | $g_{Gom} (\pm SE)$ | $g_{log} (\pm SE)$ | $RSE$ |
| --- | --- | --- | --- | --- | --- | --- | --- | --- | --- | --- |
| *Observed data* |  |  |  |  |  |  |  |  |  |  |
| VBGF | 10 | 81.07 | 0 | 0.35 | 366.3 ($\pm 207.8$) | 61.7 ($\pm 7.4$) | 0.05 ($\pm 0.04$) | - | - | 9.14 |
| Logistic | 10 | 81.28 | 0.2 | 0.31 | 217.8 ($\pm 25.6$) | 63.7 ($\pm 6.7$) | - | - | 0.24 ($\pm 0.05$) | 9.23 |
| Gompertz | 10 | 81.12 | 0.05 | 0.34 | 248.0 ($\pm 47.9$) | 62.8 ($\pm 7.0$) | - | 0.14 ($\pm 0.04$) | - | 9.17 |
| Model average | - | - | - | - | 279.7 ($\pm 118.7$) | 62.7 ($\pm 7.1$) | - | - | - | - |
| *Back-calculated* |  |  |  |  |  |  |  |  |  |  |
| VBGF | 72 | 466.7 | 0.72 | 0.33 | 369.8 ($\pm 73.4$) | 56.6 ($\pm 1.5$) | 0.05 ($\pm 0.01$) | - | - | 5.96 |
| Logistic | 72 | 467.8 | 1.8 | 0.19 | 200.9 ($\pm 7.7$) | 58.7 ($\pm 1.3$) | - | - | 0.27 ($\pm 0.02$) | 6.00 |
| Gompertz | 72 | 466.0 | 0 | 0.48 | 230.8 ($\pm 14.0$) | 57.6 ($\pm 1.4$) | - | 0.16 ($\pm 0.01$) | - | 5.93 |
| Model average | - | - | - | - | 271.1 ($\pm 74.7$) | 57.5 ($\pm 1.6$) | - | - | - | - |

$n$ is the sample size, $AIC_C$ is the small-sample bias adjusted form of Akaike's Information Criterion, $\Delta$ is the difference in $AIC_C$ values between models, $w$ are the $AIC_C$ weights, $L_\infty$ is asymptotic length parameter in cm, $L_0$ is the length-at-birth parameter in cm, $k$ is the growth completion parameter in yr<sup>-1</sup> for the VBGF, $g$ is the growth parameter for logistic and Gompertz functions (but is incomparable between the two), $SE$ is the standard error of the adjacent parameter, and $RSE$ is the residual standard error of the candidate model.

The back-calculated data provided similar growth parameters to the observed data which produced a curve with slightly smaller length-at-ages (Fig. 4). The *L*_*0*_ estimates were similar between all three candidate models and were slightly smaller than the back-calculated data (Table 2). The VBGF again provided the largest *L*_*∞*_ estimate while the logistic and Gompertz models provided estimates that were more biologically plausible (Table 2). However, for the back-calculated data the logistic model estimated a lower *L*_*∞*_ that was approximately 60 cm smaller than the estimated maximum length (∼260 cm TL; White et al. 2015). Subsequently, the logistic model had the lowest *w*_*i*_ of any candidate model for the back-calculated data (Table 2). Overall, the parameter standard errors and model residual standard errors were lower for the back-calculated data due to the addition of interpolated data (Table 2). Therefore, given that parameters for the observed and back-calculated data were similar – the back-calculated data were considered more precise. The model averaged parameters for the back-calculated data were 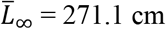 TL and 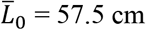 TL (Table 2). The lack of mature individuals in the sample means that this analysis should be considered as a partial growth curve with length-at-age estimates that are valid over the available age range (Table 3).

**Table 3.** Model averaged total length-at-age estimates from the back-calculated data for *Glyphis glyphis* over the age range available. 95% confidence intervals are in parentheses.

| Age | Model averaged total length estimate (cm) |
| --- | --- |
| 0 | 57 (55–60) |
| 1 | 71 (62–81) |
| 2 | 84 (69–102) |
| 3 | 98 (77–122) |
| 4 | 111 (84–141) |
| 5 | 123 (91–160) |
| 6 | 135 (99–177) |
| 7 | 146 (105–192) |
| 8 | 156 (112–206) |
| 9 | 166 (118–219) |
| 10 | 174 (123–231) |
| 11 | 182 (133–250) |

**Figure 4.**
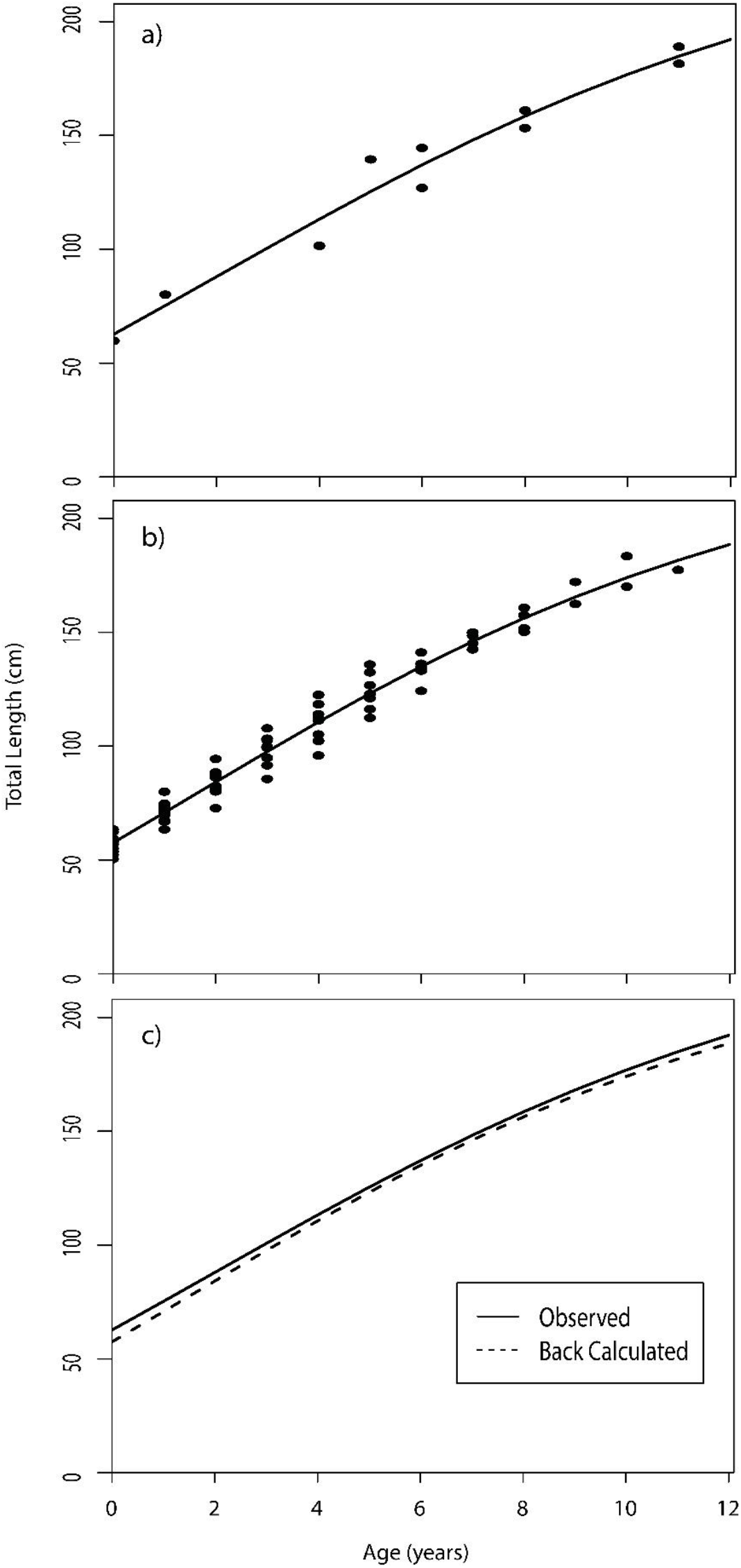
Length-at-age curves from a) observed data, b) back-calculated data, and c) a comparison of these two curves for *Glyphis glyphis* with sexes combined. Both length-at-age curves were produced from MMI results as candidate models had equal *w* for both data sets.

## Discussion

We use the smallest known sample size to generate growth curves in a shark species and provide the first age and growth estimates for any river shark species (Carcharhinidae; *Glyphis*). Both river species (*G. glyphis* and *G. garricki*) occurring in northern Australian riverine and estuarine waters are rare, threatened, and protected species precluding opportunities to acquire a large or even moderate sample size to undertake ageing through examination of vertebral band pairs. We directly targeted a very small number (n = 10) of sharks of selected size classes and applied back-calculation techniques to compliment observed band counts. Very small sample sizes have been used to produce age-at-length estimates for only a handful of rare and threatened species: Winghead Shark *Eusphyra blochii* (n = 14; Smart et al. 2013), Fossil Shark *Hemipristis elongata* (n = 14; Smart et al. 2013), and Maugean Skate *Dipturus maugeanus* (n = 13; Awruch et al. 2021).

All samples examined in this study were from juveniles or sub-adults. Adults are unknown in the Northern Territory with only a few records from small-scale fishers in Papua New Guinea (White et al. 2016). The very small sample size and the lack of mature individuals means that the results presented here should be considered a partial growth curve with length-at-age estimates that are valid over the available age range. We also caution that due to the limitations of the approach undertaken here, we were unable to validate annual growth band deposition and assumed that band pair deposition was annual. It is noted that band pair deposition can be linked to somatic growth and that counting vertebral band pairs has resulted in the underestimation of age in sharks (Harry 2018, Natanson et al. 2018).

Despite these limitations, the growth curves and parameters are biologically plausible. Given adult *G. glyphis* has never been recorded in the Northern Territory, there is not likely to be an opportunity to incorporate samples from larger size classes into the analysis. The highest age produced from this study was 11 years for a female subadult shark. We therefore assume that age-at-maturity may be in the order of 12 years. This suggests that *G. glyphis* is a relatively long-lived shark. Ageing studies on other carcharhinid species with a similar age-at-maturity have produced maximum ages of 19–30 years e.g., Grey Reef Shark *Carcharhinus amblyrhynchos* (age-at-maturity, 11 years; maximum age, 19 years; Robbins 2006) and Pigeye Shark *C. amboinensis* (age-at-maturity, 13 years; maximum age, >30 years; Tillett et al. 2011). The relatively late age-at-maturity and predicted longevity of *G. glyphis* suggests the species exhibits the conservative life-history parameters that are characteristic of large sharks and which limit their ability to recover rapidly from population depletion.

## Acknowledgements

This work was supported by the Marine Biodiversity Hub, a collaborative partnership supported through funding from the Australian Government’s National Environmental Science Program (NESP). We thank Russ Bradford, Christy Davies, Brittany Finucci, and Micha Jackson for assistance in the field, and Nic Bax, Paul Hedge, Annabel Ozimec, Roanne Ramsey, and Thor Saunders for project support.

## References

Akaike, H. (1973). Information theory as an extension of the maximum likelihood. Second International Symposium on Information Theory, Akademiai Kiado, Budapest.

Awruch, C. A., Bell, J. D., Semmens, J. M. and Lyle, J. M. (2021). Life history traits and conservation actions for the Maugean skate (Zearaja maugeana), and endangered species occupying an anthropogenically impacted estuary. Aquatic Conservation: Marine and Freshwater Ecosystems 31: 2178–2192.

Bowker, A. H. (1948). A test for symmetry in contingency tables. Journal of the American Statistical Association 43: 572–574.

Burnham, K. P. and Anderson, D. R. (2001). Kullback-Leibler information as a basis for strong inference in ecological studies. Wildlife Research 28: 111–119.

Cailliet, G. M. and Goldman, K. J. (2004). Age determination and validation in chondrichthyan fishes. In “Biology of Sharks and Their Relatives”. eds: J. Musick, J. C. Carrier and M. R. Heithaus, Boca Raton FL., CRC Press: 399–447.

Cailliet, G. M., Smith, W. D. Mollet, H. F. and Goldman, K. J. (2006). Age and growth studies of chondrichthyan fishes: the need for consistency in terminology, verification, validation, and growth function fitting. Environmental Biology of Fishes 77 (3–4): 211–228.

Campana, S. E. (1990). How reliable are growth back-calculations based on otoliths? Canadian Journal of Fisheries and Aquatic Sciences 47 (11): 2219–2227.

Campana, S. E., Annand, M. C. and McMillan, J. I. (1995). Graphical and statistical methods for determining the consistency of age determinations. Transactions of the American Fisheries Society 124 (1): 131–138.

Carlander, K. D. (1969). Handbook of Freshwater Fishery Biology, Iowa University Press, Ames.

Chang, W. Y. B. (1982). A statistical method for evaluating the reproducibility of age determination. Canadian Journal of Fisheries and Aquatic Sciences 39 (8): 1208–1210.

Evans, G. T. and Hoenig, J. M. (1998). Testing and viewing symmetry in contingency tables, with application to readers of fish ages. Biometrics 54 (2): 620–629.

Feutry, P., Berry, O., Kyne, P. M., Pillans, R. D., Hillary, R. M., Grewe, P. M., Marthick, J. R., Johnson, G., Gunasekera, R. M., Bax, N. J., Bravington, M. (2017). Inferring contemporary and historical genetic connectivity from juveniles. Molecular Ecology 26: 444–456.

Feutry, P., Devloo-Delva, F., Tran Lu y, A., Mona, S., Gunasekera, R. M., Johnson, G., Pillans, R. D., Jaccoud, D., Kilian, A., Morgan, D. L., Saunders, T., Bax, N. J. and Kyne, P. M. (2020). One panel to rule them all: DArTcap genotyping for population structure, historical demography, and kinship analyses, and its application to a threatened shark. Molecular Ecology Resources 20: 1470–1485.

Francis, R. I. C. C. (1990). Back calculation of fish length - a critical review. Journal of Fish Biology 36 (6): 883–902.

Grant, M. I., Kyne, P. M., Simpfendorfer, C. A., White, W. T., Chin, A. (2019). Categorising use patterns of non-marine environments by elasmobranchs and a review of their extinction risk. Reviews in Fish Biology and Fisheries 29: 689–710.

Harry, A. V. (2018). Evidence for systematic age underestimation in shark and ray ageing studies. Fish and Fisheries 19: 185–200.

Heupel, M. R. and Simpfendorfer, C. A. (2010). Science or slaughter: Need for lethal sampling of sharks. Conservation Biology 24: 1212–1218.

IUCN (2022). The IUCN Red List of Threatened Species. Version 2022-1. <https://www.iucnredlist.org>

IUCN Standards and Petitions Committee. (2019). Guidelines for using the IUCN Red List Categories and Criteria. Version 14. Prepared by the Standards and Petitions Committee. IUCN, Gland, Switzerland and Cambridge, UK.

Katsanevakis, S. (2006). Modelling fish growth: multi-model inference and model selection uncertainty. Fisheries Research 81: 229–235.

Katsanevakis, S. and Maravelias, C. D. (2008). Modelling fish growth: multi-model inference as a better alternative to a priori using von Bertalanffy equation. Fish and Fisheries 9 (2): 178–187.

Kyne, P. M. and Lucifora, L. O. (2022). Freshwater and euryhaline elasmobranchs. Pp. 567–602. In: Carrier, J. C., Simpfendorfer, C. A., Heithaus, M. R. and Yopak, K. E. (Eds). Biology of Sharks and Their Relatives, Third Edition. CRC Press, Boca Raton.

Kyne, P. M., Davies, C.-L., Devloo-Delva, F., Johnson, G., Amepou, Y., Grant, M. I., Green, A., Gunasekara, R. M., Harry, A. V., Lemon, T., Lindsay, R., Maloney, T., Marthick, J., Pillans, R. D., Saunders, T., Shields, A., Shields, M. and Feutry, P. (2021). Molecular analysis of newly-discovered geographic range of the threatened river shark Glyphis glyphis reveals distinct populations. Report to the National Environmental Science Program, Marine Biodiversity Hub. Charles Darwin University, Darwin and CSIRO, Hobart.

Musick, J. A. (1999). Ecology and conservation of long-lived marine animals. American Fisheries Society Symposium 23:1–10.

Natanson, L. J., Skomal, G. B., Hoffman, S. L., Porter, M. E., Goldman, K. J. and Serra, D. (2018). Age and growth of sharks: do vertebral band pairs record age? Marine and Freshwater Research 69: 1440–1452.

Pillans, R. D., Stevens, J. D., Kyne, P. M. and Salini, J. (2009). Observations on the distribution, biology, short-term movements and habitat requirements of river sharks Glyphis spp. in northern Australia. Endangered Species Research 10: 321–332.

R Core Team (2013). R: A language and environment for statistical computing. R. F. S. Computing. Vienna, Austria, R Foundation Statistical Computing.

Ricker, W. E. (1975). Computation and interpretation of biological statistics of fish populations. Bulletin of the Fisheries Research Board of Canada 191: 1–382.

Ricker, W. E. (1979). Growth rates and models. In Fish Physiology. eds: W. S. Hoar, D. J. Randall and J. R. Brett, New York NY, Academic Press. 8: 677–743.

Robbins, W. D. (2006). Abundance, demography and population structure of the grey reef shark (Carcharhinus amblyrhynchos) and the whitetip reef shark (Triaenodon obesus) (fam. Carcharhinidae). PhD Thesis. James Cook University, Townsville, Australia.

Smart, J. J., Chin, A., Baje, L., Tobin, A. J., Simpfendorfer, C. A. and White, W. T. (2017). Life history of the silvertip shark Carcharhinus albimarginatus from Papua New Guinea. Coral Reefs 36: 577–588.

Smart, J. J., Chin, A., Tobin, A. J. and Simpfendorfer, C. A. (2016). Multimodel approaches in shark and ray growth studies: strengths, weaknesses and the future. Fish and Fisheries 17: 955–971.

Smart, J. J., Harry, A. V., Tobin, A. J. and Simpfendorfer, C. A. (2013). Overcoming the constraints of low sample sizes to produce age and growth data for rare or threatened sharks. Aquatic Conservation: Marine and Freshwater Ecosystems 23 (1): 124–134.

Smith, S. E., Au, D. W. and Show, C. (1998). Intrinsic rebound potential of 26 species of Pacific sharks. Marine and Freshwater Research 49: 663–678.

Tillett, B. J., Meekan, M. G., Field, I. C., Hua, Q. and Bradshaw, C. J. A. (2011). Similar life history traits in bull (Carcharhinus leucas) and pig-eye (C. amboinensis) sharks. Marine and Freshwater Research 62: 850–860.

von Bertalanffy, L. (1938). A quantitative theory of organic growth (inquires on growth laws. II). Human Biology 10 (2): 181–213.

White, W. T., Appleyard, S. A., Sabub, B., Kyne, P. M., Harris, M., Lis, R., Baje, L., Usu, T., Smart, J. J., Corrigan, S., Yang, L. and Naylor, G. J. P. (2015). Rediscovery of the threatened river sharks, Glyphis garricki and G. glyphis, in Papua New Guinea. PLoS ONE 10 (10): e0140075.

Zhu, L., Li, L. and Liang, Z. (2009). Comparison of six statistical approaches in the selection of appropriate fish growth models. Chinese Journal of Oceanology and Limnology 27 (3): 457–467.

